# RNAIndel: discovering somatic coding indels from tumor RNA-Seq data

**DOI:** 10.1101/512749

**Authors:** Kohei Hagiwara, Liang Ding, Michael N. Edmonson, Stephen V. Rice, Scott Newman, Soheil Meshinchi, Rhonda E. Ries, Michael Rusch, Jinghui Zhang

## Abstract

Reliable identification of expressed somatic insertion/deletion (indels) is an unmet demand due to artifacts generated in PCR-based RNA-Seq library preparation and the lack of normal RNA-Seq data, presenting analytical challenges for discovery of somatic indels in tumor trasncriptome.

By implementing features characterized by PCR-free whole-genome and whole-exome sequencing into a machine-learning framework, we present RNAIndel, a tool for predicting somatic, germline and artifact indels from tumor RNA-Seq data alone. RNAIndel robustly predicts 87□93% of somatic indels from 235 samples with heterogeneous conditions, even recovering subclonal (VAF range 0.01–0.15) driver indels missed by targeted deep-sequencing, outperforming the current best-practice for RNA-Seq variant calling which had 57% sensitivity but with 12 times more false positives.

RNAIndel is freely available at https://github.com/stjude/RNAIndel

## 1. Introduction

Transcriptome sequencing (RNA-Seq) is a versatile platform for performing a multitude of cancer genomic analyses such as gene expression profiling, allele specific expression, alternative splicing and fusion transcript detection. However, variant identification in RNA-Seq is not a common practice due to the presence of artifacts introduced in library preparation, the intrinsic complexity of splicing, and RNA editing (Piskol *et al*., 2013). RNA-Seq data are predominantly generated from tumor-only samples as acquisition of a normal tissue with a comparable transcriptome is a rare practice. This lack of matching normal data further complicates somatic variant discovery in RNA-Seq. Despite these challenges, there are compelling reasons to explore RNA-Seq data for variant detection: 1) RNA variants are expressed, therefore more interpretable to cancer phenotype and clinical actionability than DNA variants; and 2) Some studies only analyze tumor specimen by RNA-Seq, and employing variant detection will make full use of the available data resources. Thus, successful development of RNA-Seq variant calling tools will make this platform an interpretable and cost-effective alternative to DNA-based whole-genome or whole-exome sequencing (DNA-Seq), the current standard platform for somatic variant detection.

Various single nucleotide variants (SNV) detection tools dedicated to RNA-Seq have been developed. SNPiR (Piskol *et al*., 2013) calls true RNA-Seq SNVs by hard-filtering calls in repetitive and low-quality regions, around splice sites, and at known RNA-editing sites. RVboost (Wang *et al*., 2014) is a machine-learning method to prioritize true SNVs trained on common SNPs in the input RNA-Seq data. eSNV-Detect (Tang *et al*., 2016) incorporates results generated from two mappers to confidently call expressed SNVs by removing mapping artifacts from individual mappers. Opossum (Oikkonen and Lise 2017) preprocesses RNA-Seq reads for SNV calling by splitting intron-spanning reads and removing spurious reads. By contrast, inde detection in transcriptome is more challenging and has been largely unexplored (Sun *et al*., 2017). Even in DNA-Seq, indel discovery suffers from a high false discovery rate (Fang *et al*., 2014). In RNA-Seq, in addition to the artifacts from library preparation due to polymerase chain reaction (PCR), alignment of spliced reads containing an indel can introduce mapping artifacts. Indels are less common than SNVs in the genome with the ratio 1:7 and even more so in coding regions with 1:43 (Ng *et al*., 2008). This low prevalence poses additional challenge for developing a robust indel detection method that optimizes sensitivity and specificity. In cancer studies, somatic indels should also be distinguished from germline indels. Therefore, somatic indel identification in tumor RNA-Seq can be formulated as a three-class classification problem where somatic, germline, and artifact indels must be considered.

Here, we introduce RNAIndel, a novel computational software that takes a tumor RNA-Seq BAM file as input, calls and annotates coding indels, and classifies them into somatic, germline and artifacts by supervised learning. RNAIndel was developed by using 765,475 indels collected from 330 pediatric tumor transcriptomes. To demonstrate that the model is not sensitive to the particular error profiles in the training set, we tested RNAIndel against two RNA-Seq datasets comprised of 235 samples generated by different library preparation protocols, read lengths, and sequencing platforms. Under these conditions, RNAIndel achieved 87□93% sensitivity with 7□10 somatic indels predicted from a total of 1,000□4,000 RNA-Seq indels per sample. RNAIndel is also flexibly designed to allow researchers to import data from their own variant caller rather than the internally-implemented caller. By demonstrating its robustness in somatic indel prediction, we anticipate that RNAIndel will augment the range of RNA-Seq applications and facilitate the investigation of expressed somatic variants.

## 2. Overview

The RNAIndel software (Fig. 1A) requires a RNA-Seq BAM file mapped by STAR (Dobin *et al*., 2013) as the input. Indel calling can be performed by the built-in Bambino (Edmonson *et al*., 2011) caller which was optimized for RNA-Seq indel calling, or by an external caller by supplying calls in the Variant Call Format (VCF) (Danecek *et al*., 2011). Indels are annotated using all RefSeq (Pruitt *et al*., 2005) isoforms containing coding exons; indels within a coding exon or in an intron region within 10 bases of a splice site (splice region) are considered coding indels and subjected to further analysis. For each coding indel, RNAlndel extracts reads covering the indel locus to retrieve the actual alignment pileup. This process also enables the incorporation of additional variations such as polymorphisms near the indel into the feature calculation. Matches to the dbSNP database (Build 151) (Sherry *et al*., 2001) are also annotated as a feature; equivalent indels are also considered during matching (Supplementary Fig. S1). For indels supported by ≥2 unique reads, prediction is made by classifiers specifically trained based on their size (i.e., 1-nt or >1-nt), which consist of an ensemble of random forest (Breiman, 2001) models. RNAIndel generates a VCF file where indel entries are parsimonious and left-aligned (Tan *et al*., 2015) to unify equivalent alignments of supporting reads (Supplementary Fig. S1). Predicted class and probability are reported in the VCF INFO field as well as dbSNP membership, ClinVar (Landrum *et al*., 2014) annotation, variant effect annotation and calculated feature values.

**Figure 1.**
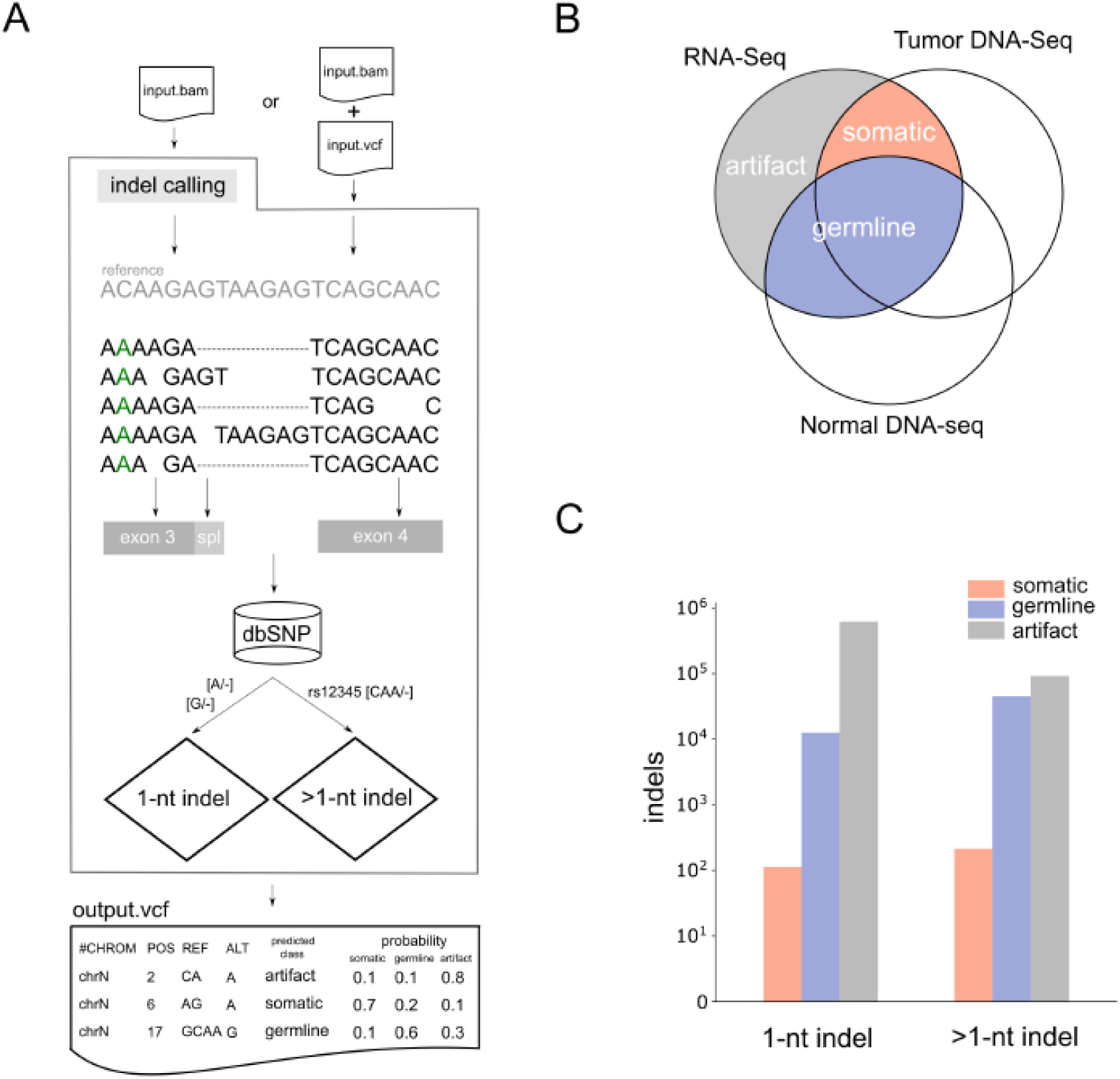
Computational framework and training dataset construction. **A**. Workflow of RNAIndel. A tumor RNA-Seq BAM file is a required input. If an optional VCF file from user’s variant caller is supplied, indel calls in the file are used for prediction. Otherwise, indel calling is performed on the input BAM file using the built-in Bambino caller. Features are calculated using alignment pileup, transcript structure, and dbSNP database. Alignments are spliced (dashed) and may contain nonreference variations which alter sequences flanking indels (C>A in green). Indels are annotated for coding exon (grey box) and splice region (light-grey box), defined as an intronic region within 10 nt of the exon boundary. After annotating the dbSNP membership, single nucleotide indels (1-nt) and multi-nucleotide indels (>1-nt) are predicted separately by random forest classifiers specifically trained for each type. Predicted class is based on the highest probability of being somatic, germine or artifact. RNAIndel outputs an annotated VCF file. **C**. Training set generated from 330 cases. Indel calls in RNA-Seq were classified somatic, germline and artifact by matching with T/N-paired WES and WGS (DNA-Seq) data. **B**. The 1-nt and >1-nt indel distribution in the categories of somatic, germline and artifacts. The class distribution of each dataset is shown in logarithm scale.

## 3. Indel Classifiers

### 3.1 Training set

Each case in the training set (N=330) was sequenced by tumor RNA-Seq, and tumor (T) / normal (N)-paired whole exome sequencing (WES) and PCR-free whole genome sequencing (WGS) (Materials and Methods). Coding indel in the training RNA-Seq dataset was labelled somatic, germline or artifact based on the paired WES and WGS analysis (Fig. 1B). Specifically, an indel in RNA-Seq was labelled somatic if it matched to a somatic indel identified by the T/N-paired DNA-Seq analysis. Expressed germline indels were defined if they were supported by the normal WGS and WES. The remaining indels, present in RNA-Seq but absent in WGS and WES, were labelled as artifacts. The resulting training set, comprised of single-nucleotide indel (1-nt) and multi-nucleotide indels (>1-nt), showed distinct distributions in the three categories of somatic, germline, and artifact (Fig. 1C) where 1-nt indels were highly enriched in artifacts. Specifically, 1-nt indels accounted for 115, 12,529, and 616,121 of somatic, germline and artifacts, respectively while >1-nt indels accounted for 213, 45,098, and 91,399 of somatic, germline and artifacts, respectively. An accompanying T/N-paired DNA-Seq analysis identified 959 somatic coding indels in this cohort, showing that 35.2% of them were expressed. Each sample harbored, on average, 0.88 ± 1.19 s.d. expressed somatic indel, ranging from 0 to 6. Top 10 frequent genes involved were *ETV6* (11 samples), *CCND3* (6), *GATA1* (6), *NOTCH1* (6), *PTCH1* (6), *BCOR* (5), *SETD2* (5), *XBP1* (5), *PTEN* (4) and *WT1* (4), highlighting the tumor-biological significance of expressed somatic indels.

### 3.2 Features

We extracted a total of 31 features from sequence/alignment, transcript/protein and database categories (Table 1 and Supplementary Methods). Several features were selected based on the strand-slippage hypothesis, a widely accepted model for explaining the mechanism by which indels are generated in the process of DNA replication (Garcia-Diaz and Kunkel, 2006). Under this model, the DNA polymerase pauses synthesis in repetitive regions, and this delay in replication allows unpaired nucleotides to transiently anneal, leading to a misaligned replication. Thus, features governing sequence complexity (feature 1□3) and annealing temperature (feature 4□7) are expected to be important parameters of this model. By contrast, insertions or deletions that are dissimilar to the flanking sequences are unlikely to be caused by strand slippage (feature 8). Cancer-associated indels may be complex (Supplementary Fig. S2) (Ye *et al*., 2016), so we define indel complexity based on misalignments near the indel site. Indel size (feature 10) is negatively correlated to prevalence, and insertions (feature 11) are rarer than deletions in the human genome (Zhang and Gerstein, 2003). Polymers of adenine or thymine are known to be more erroneous than guanine or cytosine polymers (feature 12□15) (Fang *et al*., 2014). Indels with sufficient read support are more likely to be true (feature 16□18). Mapping artifacts stem from ambiguous mapping (feature 19) or difficulty in mapping spliced reads (feature 20). Equivalent indels are alternative alignments of the identical indel sequence, a type of mapping artifact (feature 21). PCR-based genotyping can frequently create false multiallelic indels (feature 22) (Weber *et al*., 2002).

**Table 1.** Features used by RNAIndel falling into three categories. Features 1□22 characterize sequence and/or alignment properties. Features 23□30 are related to variant effects on gene products. Membership in the dbSNP database Build 151 (DB) is Feature 31. Features used before (Initial model) or after feature selection (Final model) are checked for single-nucleotide (1-nt) and multi-nucleotide indels (>1-nt). The definitions and computations of features are described in Supplementary Methods.

|  |  |  | Initial model |  | Final model |  |
| --- | --- | --- | --- | --- | --- | --- |
|  | Category | Feature | 1-nt | >1-nt | 1-nt | >1-nt |
| 1 | Sequence/Alignment | repeat | ? | ? | ? |  |
| 2 |  | lc | ? | ? |  |  |
| 3 |  | local_lc | ? | ? |  |  |
| 4 |  | gc | ? | ? |  |  |
| 5 |  | local_gc | ? | ? |  |  |
| 6 |  | strength | ? | ? |  |  |
| 7 |  | local_strength | ? | ? |  | ? |
| 8 |  | dissimilarity |  | ? |  | ? |
| 9 |  | indel_complexity | ? | ? | ? | ? |
| 10 |  | indel_size |  | ? |  | ? |
| 11 |  | is_ins | ? | ? |  | ? |
| 12 |  | is_at_ins | ? |  | ? |  |
| 13 |  | is_at_del | ? |  | ? |  |
| 14 |  | is_gc_ins | ? |  |  |  |
| 15 |  | is_gc_del | ? |  |  |  |
| 16 |  | ref_count | ? | ? | ? | ? |
| 17 |  | alt_count | ? | ? | ? | ? |
| 18 |  | is_bidirectional | ? | ? |  | ? |
| 19 |  | is_uniq_mapped | ? | ? | ? | ? |
| 20 |  | is_near_exon_boundary | ? | ? | ? | ? |
| 21 |  | equivalence_exists | ? | ? |  |  |
| 22 |  | is_multiallelic | ? | ? |  | ? |
| 23 | Transcript/Protein | is_inframe |  | ? |  |  |
| 24 |  | is_splice | ? | ? |  |  |
| 25 |  | is_truncating | ? | ? |  | ? |
| 26 |  | is_in_cdd |  | ? |  |  |
| 27 |  | indel_location | ? | ? |  |  |
| 28 |  | is_nmd_insensitive | ? | ? | ? |  |
| 29 |  | indels_per_gene | ? | ? | ? | ? |
| 30 |  | coding_seq_length | ? | ? |  |  |
| 31 | DB | is_on_dbsnp | ? | ? | ? | ? |

Indels were also characterized in terms of variant effect. In-frame indels are non-truncating unless they create a *de novo* stop codon. Indels in splicing regions may not affect splicing unless they destroy the splice motif (feature 23□25). Further, in-frame indels may be less deleterious if they occur outside of conserved domains (feature 26). The relative location of indels (feature 27) within proteins are bimodal around the N and C-termini. It has been postulated that transcripts with N-terminual indels can be rescued by an alternative downstream start codon, while a subset of C-terminus truncations may retain the all functional domains (Ng *et al*., 2008). Indels in the first and last exons are therefore known to be less sensitive to nonsense mediated decay (NMD) (feature 28) (Popp and Maquat, 2016). We also hypothesized that the number of true indels in a gene is expected to be few. The number of detected indels in a gene were normalized by dividing by the coding sequence length (feature 29□30). Finally, indels present in dbSNP are considered more likely to be germline (feature 31).

### 3.3. Optimization

Somatic indel discovery is the most likely application for RNAIndel, a context in which sensitivity is especially critical. For this reason, models were optimized favoring true positive rate (TPR), or sensitivity, of somatic indels over false discovery rate (FDR). As the training set was particularly imbalanced towards artifacts (Fig. 1C), we downsampled the training set to somatic: germline: artifact = 1: 1: *x*. The ratio for arifact was determined to maximize *F_10_* for somatic class on a 5-fold cross-validataion (Materials and Methods). The downsampled training sets were at somatic: germline: artifact = 1: 1: 18 for 1-nt with a *F_10_* of 0.7 and 1: 1: 6 for >1-nt with a *F_10_* of 0.79 (Supplementary Table S1). The models were further refined for somatic prediction. An optimal subset of the 27 features for each indel class was selected by a greedy best-first search for somatic TPR improvement (Materials and Methods), leading to a set of 11 features for 1-nt and 14 features for >1-nt indel type, respectively (Table 1). Performance metrics for each class are summarized (Table 2). Although optimized for somatic prediction, the final models accurately predicted germline and artifact indels as well.

**Table 2.** Performance of the final model for 1-nt and >1-nt indel prediction. TPR (sensitivity): true positive rate; FPR (1-specificity): false positive rate; F_β_: generalized F-score. β=10 for somatic, 1 otherwise. Hand-Till: Hand-Till’s measure, a generalization of AUC to multi-class problem (Methods).

| Somatic | TPR | FPR | $F_{\beta}$ | Hand-Till |
| --- | --- | --- | --- | --- |
| 1-nt | 0.870 | 0.003 | 0.741 | 0.979 |
| > 1-nt | 0.897 | 0.015 | 0.818 | 0.982 |
| Germline |  |  |  |  |
| 1-nt | 0.952 | 0.001 | 0.954 | 0.985 |
| > 1-nt | 0.958 | 0.005 | 0.964 | 0.985 |
| Artifact |  |  |  |  |
| 1-nt | 0.996 | 0.031 | 0.997 | 0.993 |
| > 1-nt | 0.985 | 0.016 | 0.986 | 0.995 |

The selected features provided distinct profiles of somatic, germline and artifact indels. Somatic indels were longer and dissimilar to the flanking sequences. Complex indels were rare but highly specific to the somatic class (Fig. 2). Somatic indels appeared at any variant allele frequency (VAF) below 0.5 (Supplementary Fig. S3A). Germline indels were predominantly non-truncating or enriched for

**Figure 2.**
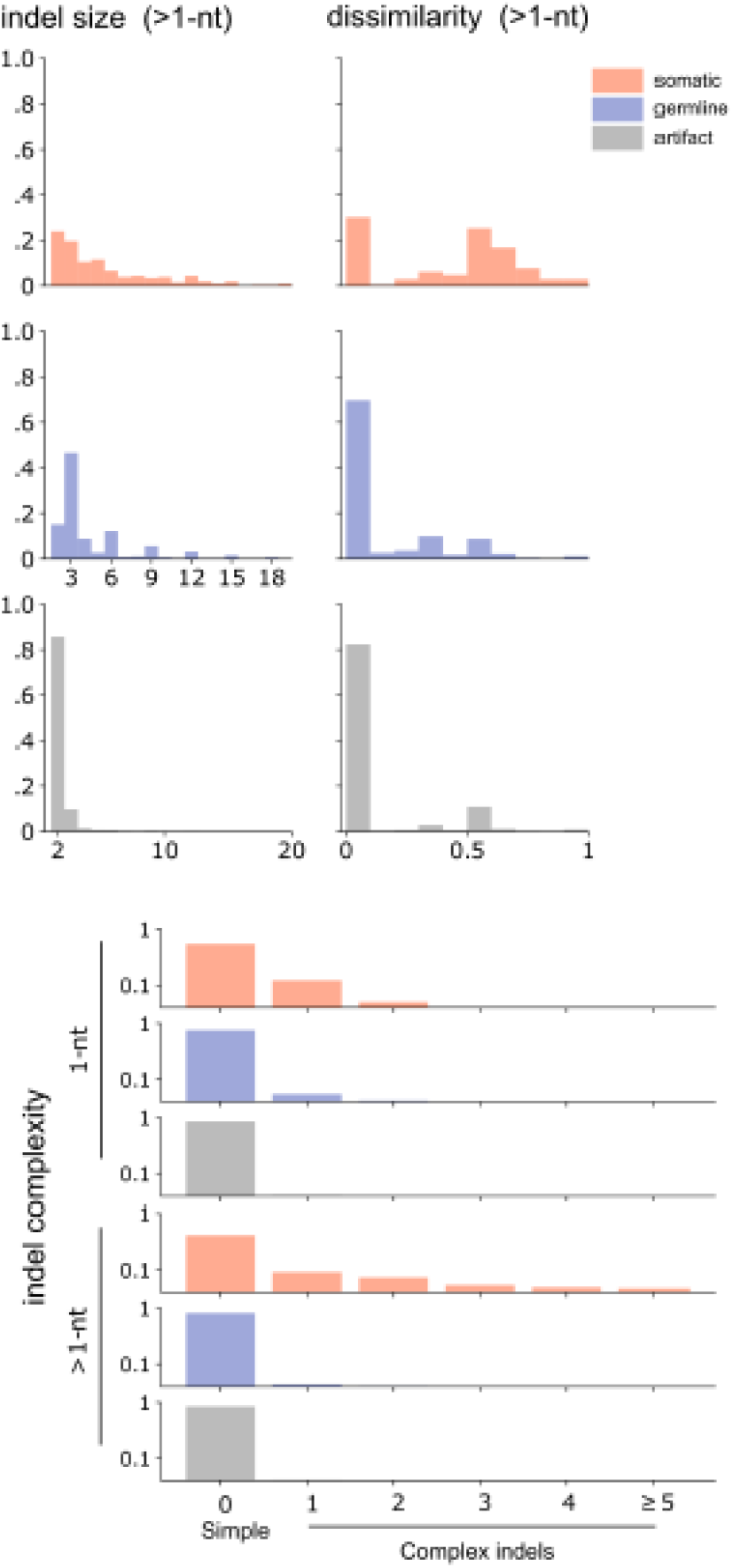
Somatic indel features. Features ‘indel size’ and ‘dissimilarity’ for >1-nt indel and ‘indel complexity’ for both 1-nt and >1-nt indels were selected in the final model. Somatic indels appeared longer and their lengths were frequently not divisible by three as opposed to germline indels. The inserted or deleted sequences in somatic indels were more dissimilar to the flanking sequences. Complex indels (Supplementary Fig. S2) were almost exclusively found in the somatic class.

NMD-insensitivity if truncating, and were mostly found in the dbSNP Build 151 database (Supplementary Fig. S3B). Artifacts were found in repetitive sequence with slightly lower nucleotide strength, a strong predictor of annealing temperature (Khandelwal and Bhyravabhotla, 2010), and were predominantly in the lowest VAF range (Supplementary Fig. S3A and S3C).

## 4. Performance

### 4.1. Somatic indel discovery from tumor RNA-Seq data

The training set was generated on an Illumina HiSeq 4000 using total RNA library with 125-nt read length. To test if the model was over-fitted to this particular specification, the trained model was evaluated using two unrelated datasets (TestSet1 and TestSet2) with varying library and sequencing specifications (Materials and Methods). Despite the technical differences between these datasets, RNAIndel was able to predict somatic indels at 87□93% TPR, performance comparable to the TPR obtained by the 5-fold CV of the training set (Table 2). Stringency was also reasonable: 3□6 indels were predicted to be somatic for 1-nt or >1-nt indel per sample (Table 3 and Supplementary Tables S2□S5). Prediction of low-frequency indels (VAF<0.15) was also sensitive with a TPR of 81% (21/26) in TestSet1 and 92% (12/13) in TestSet2. Notably, in TestSet2, five previously unknown indels with low VAF in known cancer genes were predicted as somatic by RNAIndel, and ultimately validated by manual review of targeted capture exome sequencing (Supplementary Table S6): *EP300* Y207fs (VAF: 0.1), *CEBPA* P23fs (0.17), *RAD21* D543fs (0.02), *KIT* Y418_D419>Y (0.15), and *CREBBP* S1767fs (0.01).

**Table 3.** Somatic prediction in test datasets with different conditions.

|  | Library | Read Length | Sequencer | TPR |  | Median indels predicted somatic per sample | Median indels per sample |
| --- | --- | --- | --- | --- | --- | --- | --- |
| TestSet1<br>N = 77 | Total RNA | 100 | HiSeq2000 or 2500 | 1-nt | 0.882 (15/17) | 6 | 2318 |
|  |  |  |  | >1-nt | 0.875 (35/40) | 3 | 311 |
| TestSet2<br>N = 158 | Poly A | 75 | HiSeq2000 | 1-nt | 0.909 (20/22) | 3 | 1036 |
|  |  |  |  | >1-nt | 0.934 (57/61) | 4 | 202 |

In addition to *de novo* indel analysis, RNAIndel also supports custom filtering by an optional user-defined indel panel, which is used to refine somatic prediction. Putative somatic indels matching this panel are assigned to germline or artifact, whichever has the higher probability (Fig. 3). This reclassification rule obviates the need to label non-somatic indels as germline or artifact, which can be difficult in the absence of the DNA evidence. Such a non-somatic indel panel would ideally be compiled from normal RNA-Seq data, which may not be available; however, we found that germline and artifact indels misclassified as somatic frequently recur, which offers a practical strategy for compiling an exclusion panel if users have a dataset with DNA- and RNA-Seq performed. For example, we compiled an exclusion panel by collecting non-somatic indels in the training set that were misclassified as somatic in ≥3 samples in the 5-fold CV of the final model. After applying the exclusion panel to the predictions in the two independent test sets, the median number of putative indels were 3 (1-nt) and 2 (>1-nt) for TestSet1 and 1 (1-nt and >1-nt) for TestSet2.

**Figure 3.**
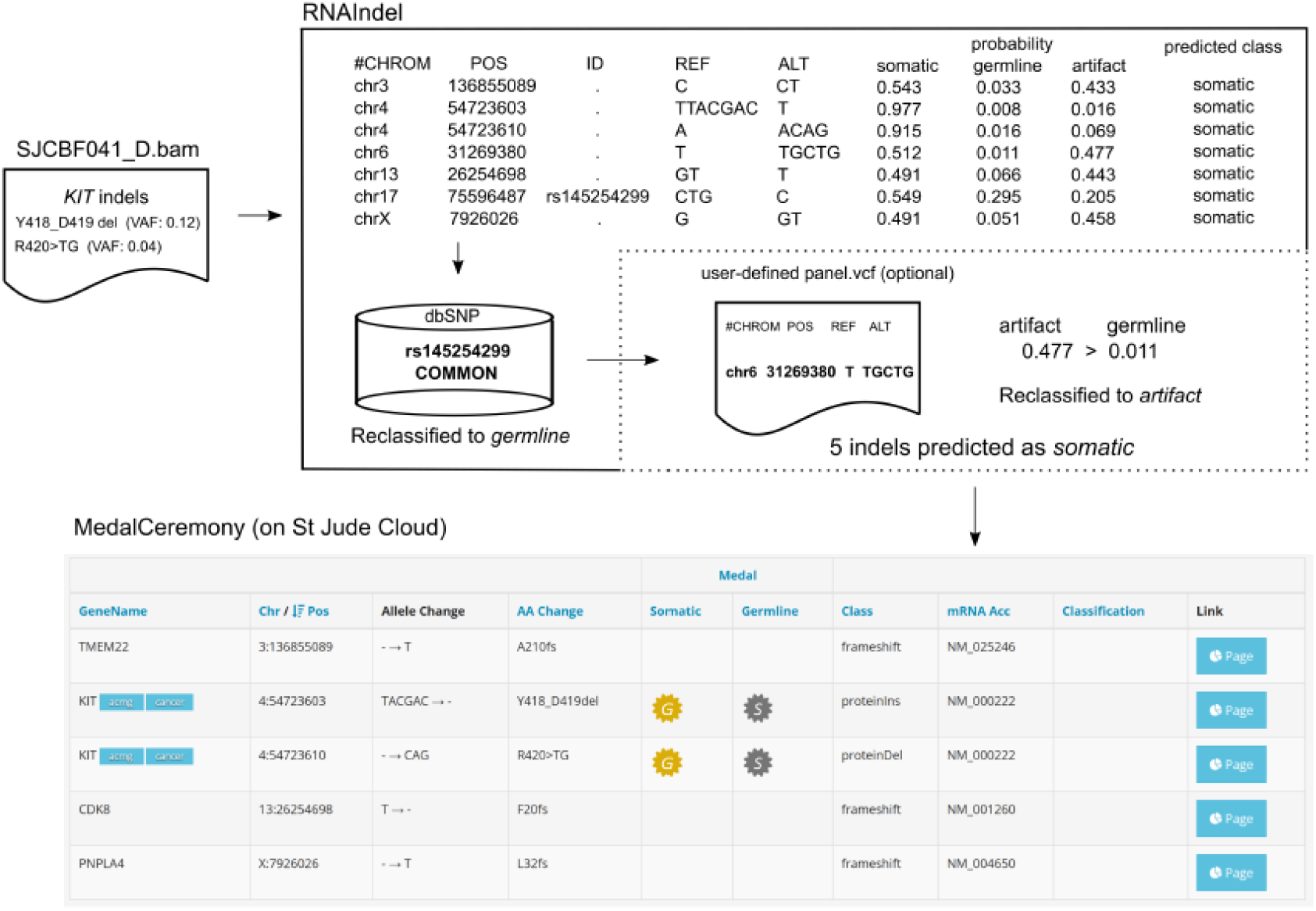
Pathogenic indel discovery from RNA-Seq. An example workflow is shown with a leukemia sample SJCBF04_D which harbors two distinct subclonal indels in the *KIT* oncogene based on WGS and WES analysis: Y418_D419del (VAF: 0.12) and R420>TG (0.04). Before reclassification, seven indels were predicted as somatic out of 3027 coding indels. The 2-nt deletion at chr17:75596487 matched rs145254299, a common indel in the dbSNP database, was reclassified as germline. Somatic prediction can be further refined by optionally supplying a user-defined exclusion panel (circumscribed in dashed line). Putative somatic indels in the panel are reclassified to germline or artifact class, whichever has the higher probability. Here, the indel on chromosome 6 matched a recurrent non-somatic indel in the panel prepared from the training set and was reclassified to artifact whose probability (0.477) was higher than germline (0.011). By uploading the output VCF file to the St Jude Cloud, the MedalCeremony algorithm prioritizes indels by pathogenicity. The *KIT* indels as somatic status were prioritized with the highest pathogenicity rank ‘gold’ (the ‘G’ symbol).

Users can also prioritize the RNAIndel prediction for pathogenicity by uploading the output VCF file to the St Jude Cloud tool PeCan PIE (https://platform.stjude.cloud/tools/pecan_pie), where the MedalCeremony algorithm (Zhang *et al*., 2015) ranks variant pathogenicity into four tiers: gold, silver, bronze, and no medal. In the example workflow (Fig. 3), the two subclonal *KIT* indels as somatic mutation were rated ‘gold’, the highest pathogenicity rank, while the other indels did not receive a medal. Details of RNAIndel’s performance on pathogenic indel prediction are presented below.

### 4.2 Working with an external variant caller— A GATK example

In addition to calling indels from a BAM file, RNAIndel can accept a VCF file made by an external variant caller (Fig. 1A). We chose GATK-HC to illustrate this feature for two reasons. First, GATK Best Practice of RNA-Seq variant calling has been documented in detail for GATK-HC with STAR (Materials and Methods). Second, the STAR/GATK-HC pipeline performed the best in a recent comparison study where combinations of a RNA-Seq mapper and a variant caller were tested for detecting known somatic *EGFR* indels in lung cancer (Sun *et al*., 2017). Following this procedure, we evaluated the performance of three approaches for detecting expressed pathogenic indels: RNAIndel with the built-in caller, RNAIndel with GATK-HC, and the Best Practice-based approach without RNAIndel (Fig. 4A). We used TestSet1 which contained 23 pathogenic indels classified by our MedalCeremony algorithm (Supplementary Table S7) and the RNA-Seq data were analyzed by the three approaches followed by the MedalCeremony classification. RNAIndel with the built-in caller achieved the highest sensitivity (22/23) with one misclassifed as artifact (Supplementary Table S7). The prediction from the combination of RNAIndel and GATK-HC had the fewest false positives (a total of 7 artifacts were predicted), but this was achieved at a cost of sensitivity: only 14 of the 23 pathogenic indels were predicted as somatic (the remaining 9 indels were not detected by GATK-HC (Supplementary Table S7)). However, for true indels predicted as somatic by this approach, the predictions coincided with those made by RNAIndel with the built-in caller (14 somatic and 3 germline), suggesting that the prediction was robust between the callers. The Best Practice prediction, i.e., the STAR/GATK-HC pipeline, was the noisiest with 442 artifacts, a rate 10 times higher than the first two approaches and also the lowest in sensitivity: only 13 of the 23 indels were correctly predicted with 1 filtered and 9 undetected (Fig. 4B, and Supplementary Table S7). While the RNAIndel-based approaches predicted 3 germline indels as somatic, two were curated as cancer predisposition mutations on ClinVar (Supplementary Table S8): *RAD50 K994_E995fs* (rs587780154) and *BRCA2 P1062_Q1063fs* (rs80359374).

**Figure 4.**
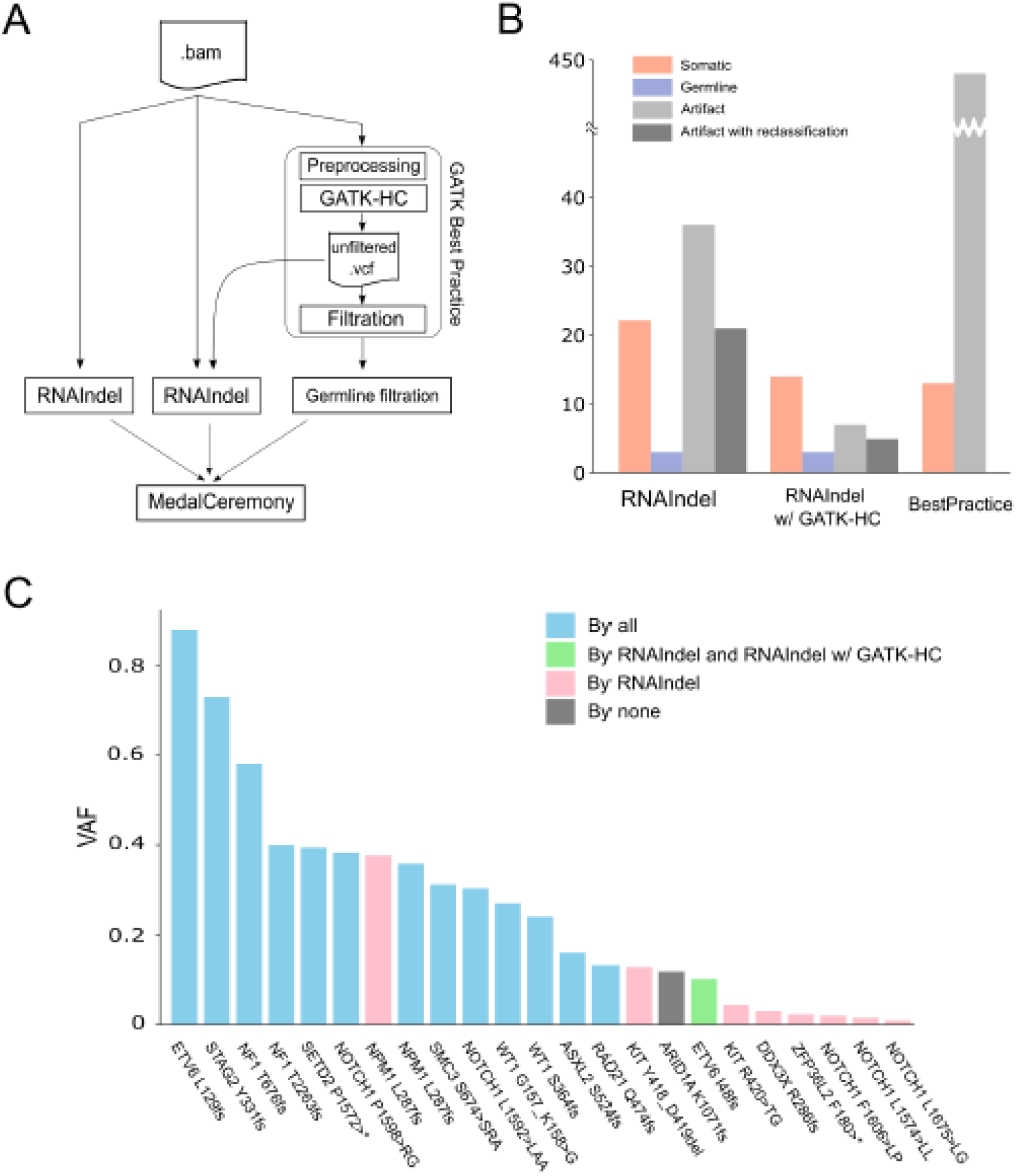
Workflow and performance using GATK as the external variant caller. RNAIndel is flexibly designed to work with an external variant caller via a user-provided VCF file. GATK-HaplotypeCaller (GATK-HC) was used as an example. **A**. TestSet1 (n=77) was screened for pathogenic somatic indels by three approaches. RNAIndel with built-in caller (left) used a RNA-Seq BAM file as input. RNAIndel with GATK-HC (middle) accepted the unfiltered VCF file from GATK-HC and the RNA-Seq BAM file as input. Best Practice-based approach (right) filters artifacts but does not discriminate somatic and germline calls. For a fair comparison, germline calls in this approach were removed by matching the normal WGS data (Germline filtration). Indels predicted as somatic by each method were prioritized by the MedalCeremony algorithm. Predicted somatic indels were defined pathogenic if prioritized as such by MedalCeremony. **B**. Predicted pathogenic indels by these approaches are shown in the bar plot. The number of artifacts is presented without (gray) and with reclassification by the panel of non-somatic indels prepared from the training set (darkgray). **C**. Of 23 known pathogenic somatic indels in this dataset, 13 indels were detected and predicted as somatic by all approaches (light blue). The *ETV6* I48fs frameshift was detected and predicted as somatic by RNAIndel and RNAIndel with GATK-HC, but filtered by the Best Practice Filtration step (green). Eight indels with lower VAF values were detected and predicted as somatic by RNAIndel (pink), but not detected by the other approaches. The *ARID1A* K1072fs frameshift was detected but predicted as artifact by RNAIndel, and not detected by the other approaches (gray).

Because GATK-HC locally assembles haplotypes while detecting variants, it may not report indels as aligned in the BAM file, and so these may not be recovered by RNAIndel’s indel search (Methods). In TestSet1, nearly half the unrecovered indels (48.6%) were located in reads spanning intron-exon boundaries. Unrecovered exonic indels were most frequently located in human leukocyte antigen (HLA) loci (12.5%). For further investigation, 100 unrecovered indels were randomly selected and reviewed: 47 cases were in intron-exon boundaries, 14 at HLA loci and 39 non-HLA exon cases (Supplementary Table S9). Remapping to the genome by the BLAT algorithm (Kent, 2002) confirmed that 44 of the 47 intron cases were mis-alignments over splice junctions. The remaining 3 cases were ambiguous, with low quality mappings. The high degree of genetic diversity at HLA loci likely explains the unsuccessful recovery. The 39 non-HLA exonic indels were also re-mapped by BLAT, revealing 17 mis-mappings. Of the remaining 22 exonic indels, 15 were found in repeats where somatic indels are less frequent as characterized in this study, 4 were in non-repetitive regions and 3 cases should have been realigned as a SNV or a cluster of SNVs rather than an indel. Considering that a majority of these indels were mismappings or in HLA loci, the impact of the underreporting may be marginal to the somatic prediction; indeed, all of the 23 somatic indels were successfully recovered in TestSet1.

## 5. Discussion

We developed RNAIndel, a machine-learning based method to classify coding indel calls in RNA-Seq data. The performance of a machine learning method is constrained by the quality of the training set. In our initial assessment, nearly one third of RNA-Seq indels supported by WES were not validated by further investigations, suggesting that they were PCR artifacts common to both platforms. To construct a high-quality training set, we employed three-platform clinical sequencing of WGS, WES and RNA-Seq with WGS data generated from a PCR-free library protocol (Rusch *et al*., 2018). Previously, we have shown that this approach achieved a positive predictive rate of 98.8% and a sensitivity of 94.3% for exonic indels, assuring the quality of the training set (Rusch *et al*., 2018). A possible limitation of the training set is that indels in the set were predominantly small indels with the maximum length being 23-nt. Analysis of larger indels may require tools designed for structural variation detection. This method was optimized and evaluated for somatic indel prediction using two unrelated test datasets generated under varying conditions. RNAIndel robustly predicted somatic indels with a TPR of ~0.9. The ability to predict low-frequency variants is particularly important as tumor heterogeneity is the source of tumor clonal evolution causing resistance to cancer treatment. However, false negatives are frequent in these low VAF variants because artifacts are predominantly distributed in this VAF range (Supplementary Fig. S3A). RNAIndel is able to discern subclonal indels with VAF <0.15 from artifacts at a TPR >0.8. This sensitivity enabled the discovery of novel somatic indels in TestSet2 which had not been identified by 500× targeted sequencing, possibly due to subclonality.

Features assigned to variants by the software are used to inform its classification logic and at times may unveil interesting biological insights. For example, the strand-slippage model predicts a simple indel with the flanking sequence. Two of our assigned features capture this state; the feature ‘dissimilarity’ is set to zero if the indel matches its flanking sequence, and the feature ‘indel_complexity’ is set to zero for simple indels. Interestingly, somatic indels frequently deviated from the slippage model, while germline and artifact indels were generally consistent with it (Fig. 2). This may be because the mechanism of somatic indel acquisition differs from that of germline due to the instability of cancer genome. Alternatively, one may speculate that a high proportion of indels compatible with the slippage model tend to be under weaker selection pressure and may hence appear in germline. For example, a triplet indel in a tandem tri-nucleotide repeat region is a pattern typical of germline indels and can be explained by strand slippage. This type of indel may have limited impact on protein function due to the possible redundancy of the repetitive amino acid molecules.

Like other variant discovery tools, RNAIndel does expect users to review the predicted outputs. In our review, single nucleotide indels (1-nt) with ‘alt_count’ being 2 or 3 appeared as a major source of misclassification. As a strategy to review such indels, we recommend applying a panel of non-somatic indels (Fig. 3), gene prioritization (Fig. 3), using another indel caller (Fig. 4) to confirm consensus, visual inspection of the indel alignments, and reasoning by user’s domain knowledge of the disease being investigated. Despite this limitation, RNAIndel enables an unbiased screening of somatic indels from tumor RNA-Seq data alone; an application of RNA-Seq data which has not been attempted due to the lack of suitable tools.

## 6. Materials and Methods

### 6.1. Datasets

All RNA-Seq data sets in this study, which consisted of a training set and two test sets, were mapped by STAR in 2-pass mode to GRCh38 (Supplementary Methods).

The training set was a dataset comprised of paired tumor (T) / normal (N) whole WGS and WES, and tumor RNA-Seq generated from 330 pediatric cancer patients from 17 major cancer types (Supplementary Table S10). Importantly, the WGS libraries were prepared by a PCR-free protocol to minimize PCR-artifacts. The details of nucleic acid extraction, library preparation, sequencing and variant detection were previously described (Rusch *et al*., 2018).

Two public datasets were used as test sets. The first set (TestSet1) was comprised of 77 RNA-Seq samples of 20 tumor types (Supplementary Table S2) previously used for developing a clinical pipeline (Rusch *et al*., 2018) (St Jude Cloud data accession SJC-DS-1003 or EGA accession EGAS00001002217). Total RNA-Seq libraries were sequenced on Illumina HiSeq2000 or 2500 with 100-nt read length. The ground-truth somatic indels in TestSet1 were based on paired T/N DNA-Seq analysis performed on the samples (Supplementary Table S3). The second test set (TestSet2) included 158 acute myeloid leukaemia samples by NCI TARGET project (https://ocg.cancer.gov/programs/target) (Supplementary Table S4). The RNA-Seq data (dbGaP study identifier phs000465) were generated at Canada’s Michael Smith Genome Sciences Centre (BCCA-GSC) using strand-specific polyA+ RNA-Seq library preparation protocol and an Illumina HiSeq2000 sequencer with 75-nt read length. The ground-truth somatic indels in TestSet2 were compiled from the paired-T/N WGS analysis by the Complete Genomics, Inc. (CGI) Cancer Sequencing service pipeline version 2 (Ma *et al*., 2018) and targeted capture sequencing analysis (Bolouri *et al*., 2018) by Strelka (Saunders *et al*., 2018) (Supplementary Table S5).

### 6.2. Performance metrics

As performance metrics for overall classification are sensitive to class imbalance, we used class-specific metrics for evaluation. For class *i*, true positive (TP), true negative (TN), false positive (FP) and false negative (FN) are defined by:

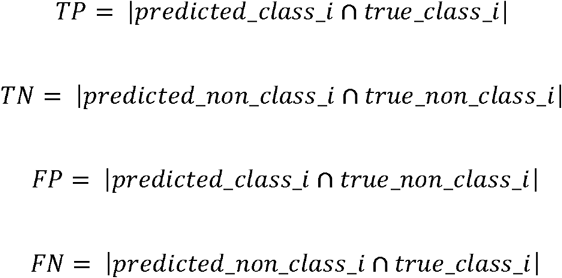

-where | | denotes the set size. True positive rate (TPR), false positive rate (FPR), and false discovery rate (FDR) are defined by TP/(TP+FN), FP/(FP+TN), and FP/(FP+TP), respectively.

A generalized F-score (*F_β_*) is formulated as:

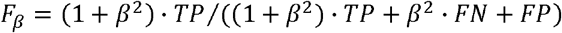

-where *β* is a weighting parameter for TP. The Hand-Till measure (M) is a generalization of the area under the curve (AUC) to multi-class classification (Hand and Till, 2001). This quantifies the separation of classes in terms of prediction probability. For three classes *i, j* and *k*, the Hand-Till measure for class is defined:

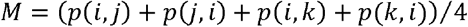

-where *p*(*x, y*) is the probability that a randomly chosen instance from class *x* has a higher prediction probability of being in class *x* than a randomly chosen class *y* instance’s probability of being class *x*.

### 6.3. Cross validation and Feature selection

Models were evaluated in the actual imbalanced distribution by 5-fold cross-validation (5-fold CV). In the first fold, 20% of the data was held out unsampled as a validation set and the training set was downsampled from the remaining 80%. Trained models were evaluated using the unsampled validation set. This process was repeated to the fifth fold by rotating the validation set portion.

Features were selected in a greedy best-first search by maximizing the true positive rate (TPR) for somatic indel. The search procedure began with an empty set of selected features (selected set) and the set of all features (feature set). In the first iteration, the model with each feature was separately evaluated in 5-fold CV. The feature that achieved the maximum TPR was added to the selected set and removed from the feature set. If features tied for the highest TPR, the feature with the minimum false discovery rate (FDR) was chosen. The next iteration examined the combination of the selected feature and one of the remaining features. The highest TPR feature was similarly added to the selected set and removed from the feature set. This procedure was continued while the model’s TPR increased.

### 6.4 RNA-Seq variant calling by GATK

Variant calling of BAM files were performed by HaplotypeCaller in the Genome Analysis Toolkit ver. 4.0.2.1. (GATK-HC) (DePristo *et al*., 2011). The workflow closely followed the GATK Best Practice for calling variants in RNA-Seq (https://software.broadinstitute.org/gatk/documentation/article.php?id=3891). The Best Practice protocol filters spurious calls but does not distinguish somatic and germline calls. For a fair comparison with RNAIndel, which distinguishes somatic, germline and artifact calls, germline calls in the Best Practice protocol were filtered by matching the normal DNA-Seq data.

### 6.5. Indel recovery

RNAIndel uses actual indel alignments in the BAM file for feature calculation. For this calculation, indels reported from the caller are searched in the BAM file. RNAIndel first searches for indels equivalent to the reported indel and merges them (Supplementary Fig. S1). If no equivalent indels are found, the nearest indel from the reported locus within a window of ±5-nt is used for analysis as proxy. These substituted cases are labelled as such in the output VCF file. Typically, 100% of indels reported from the built-in Bambino caller are successfully recovered. When GATK-HC is used as an external caller, ~90% of indels are recovered; additional discussion can be found below (Results).

## Supplementary materials

Supplementary Figures S1□S3 in SupplementaryFigures.docx

Supplementary Tables S1□S10 in SupplementaryData.xlsx

SupplementaryMethods.docx

