## SupplementaryMethod for "RNAIndel: discovering somatic coding indels from tumor RNA-Seq data"

**STAR parameters**

GTF file: gencode.v27.annotation.gtf

--outSAMunmapped Within

--outSAMstrandField intronMotif

--outSAMtype SAM

--outSAMattributes NH HI AS nM NM MD XS

--outFilterMultimapScoreRange 1

--outFilterMultimapNmax 20

--outFilterMismatchNmax 10

--alignIntronMax 500000

--alignMatesGapMax 1000000

--sjdbScore 2

--alignSJDBoverhangMin 1

--outFilterMatchNminOverLread 0.66

--outFilterScoreMinOverLread 0.66

--limitSjdbInsertNsj 2000000

--twopassMode Basic"

**Feature description**

1. ‘repeat’

The number of indel sequence repeats in the flanking sequences.

2. ‘lc’

The linguistic complexity (LC) in the flanking 50-nt reference sequence.

LC quantifies the diversity of *k*-mers in a nucleotide sequence. For *n*-nt DNA sequence, the *k*-mer usage ($U_{k}$) is defined by a ratio of the observed number of *k*-mers ($O_{k}$) and the maximal number of different *k*-mers that a *n*-bp sequence can contain (Trifonov 1990, Gabrielian and Bolshoy 1999):

$$U_{k}={O_{k}}/{min\left( 4^{k}, n-k+1 \right)}.$$

The LC value is given as the product over *1* to *n*:

$$LC= \prod_{1 \leq k \leq n} U_{k}.$$

3. ‘local_lc’

The linguistic complexity in the flanking 6-nt read sequence. The smaller linguistic complexity of the 5’ and 3’ flanking sequences is used as feature.

4. ‘gc’

The GC content in the flanking 50-nt reference sequence.

5. ‘local_gc’

The GC content in the flanking 6-nt read sequence.

6. ‘strength’

The DNA strength in the flanking 50-nt reference sequence.

DNA strength quantifies the thermodynamic stability of the sequence by considering the contribution from hydrogen bonding and π–π electron interaction (stacking). The DNA strength parameter reported in (Table 1 in Khandelwal and Bhyravabhotla 2010) is augmented to accommodate ‘N’ in addition to ‘A’, ‘T’, ‘C’, and ‘G’, by averaging possible combinations. For example, the value for ‘AN’ is the average of ‘AA’ (5), ‘AT’ (7), ‘AC’ (10) and ‘AG’ (8). For *n*-nt DNA sequence ($s$), the strength value is defined:

$${\sum_{1 \leq i \leq n-1} strength\_parameter(s_{i}s_{i+1})}/n$$

Augmented DNA strength parameters:

| GC: 13 | CC: 11 | GG: 11 | CG: 10 | AC: 10 | TC: 8 | AG: 8 | TG: 7 |
| --- | --- | --- | --- | --- | --- | --- | --- |
| GT: 10 | CT: 8 | GA: 8 | CA: 7 | AT: 7 | TT: 5 | AA: 5 | TA: 4 |
| AN: 7.5 | CN: 9 | GN: 10.5 | TN: 6 | NA: 6 | NC: 10.5 | NG: 9 | NT: 7.5 |
| NN: 8.25 |  |  |  |  |  |  |  |

7. ‘local_strength’

The DNA strength in the flanking 6-nt read sequence.

8. ‘dissimilarity’

The smaller edit distance with unit cost between the indel sequence and flanking read sequence. For *n*-nt indel sequence ($idl\_seq$), dissimilarity is defined:

$$min(edit\_distance(idl\_seq, 5\_flk),edit\_distance(idl\_seq,3\_flk))$$

where $5\_flk$ and $3\_flk$ are 5’ and 3’ *n*-nt flanking sequences.

9. ‘indel_complexity’

The edit distance between the flanking indel read and non-indel read sequences. The minimum over all combinations is used as a feature.

For a combination of indel and non-indel reads, the complexity is defined:

$$edit\_distance(5'\_flk\_of\_indel,5'\_flk\_of\_non\_indel)+ edit\_distance(3'\_flk\_of\_indel,3'\_flk\_of\_non\_indel)$$

where $5^{'}\_flk\_of\_indel$ and $3^{'}\_flk\_of\_indel$ are 6-nt flanking indel read sequences and $5^{'}\_flk\_of\_non\_indel$ and $3^{'}\_flk\_of\_non\_indel$ are 6-nt flanking non-indel read sequences

To consider cases where germline polymorphisms exist near the indel (Fig. S2), the complexity is calculated for all indel and non-indel combinations and the minimum value is used as feature.

10. ‘indel_size’

The length of the indel sequence.

11. ‘is_ins’

True for insertion, False for deletion.

12. ‘is_at_ins’

True for insertion of ‘A’ or ‘T’, False otherwise.

13. ‘is_at_del’

True for deletion of ‘A’ or ‘T’, False otherwise.

14. ‘is_gc_ins’

True for insertion of ‘G’ or ‘C’, False otherwise.

15. ‘is_gc_del’

True for deletion of ‘G’ or ‘C’, False otherwise.

17. ‘ref_count’

The number of unique reads supporting non-indel sequences.

18. ‘alt_count’

The number of unique reads supporting the indel sequence.

19. ‘is_uniq_mapped’

True if uniquely mapped by STAR, False otherwise.

20. ‘is_near_boundary’

True if the indel is exonic and within 2-nt to the exon boundary, False otherwise.

21. ‘equivalent_exists’

True if equivalent alignments are detected for the indel, False otherwise.

22. ‘is_multiallelic’

True if more than one indel sequences are detected at the same locus, False otherwise.

23. ‘is_inframe’

True if three conditions are satisfied, False otherwise: 1) annotated in coding exon, 2) the indel length is a multiple of three, and 3) the indel does not create a *de novo* stop codon.

24. ‘is_splice’

True if the indel is intronic and within 10-nt to the exon boundary, False otherwise.

25. ‘is_truncating’

True if either of the conditions are satisfied, False otherwise: 1) annotated in coding exon and its length is not a multiple of three, 2) annotated in coding exon, its length is a multiple of three, and creates a *de novo* stop codon, 3) True for ‘is_splice’ and the indel destroys the canonical splicing motif GT-AG.

26. ‘is_in_cdd’

True if the indel is annotated in a domain registered in the Conserved Domain Database (<https://www.ncbi.nlm.nih.gov/Structure/cdd/cdd.shtml>) (Marchler-Bauer et al.), False otherwise.

27. ‘indel_location’

The relative location of the indel within the coding sequence. The median value is used as a feature if the indel is annotated for multiple isoforms.

28. ‘is_nmd_insensitive’

True if the indel is located in the first or last coding exon, False otherwise.

29. ‘indels_per_gene’

The number of detected indels belonging to a gene normalized by the kilo-coding sequence length. For example, when 10 indels are detected in a gene whose coding sequence is 2000-nt long, the value is 5.0 (= 10 indels / 2 kilo). The value is averaged over isoforms if the gene has multiple isoforms.

30. ‘coding_seq_length’

The length of coding exons. The median value is used as a feature if the indel is annotated for multiple isoforms.

31. ‘is_on_dbsnp’

True if at least one equivalent indel is found in the dbSNP database, False otherwise.
