## SupplementaryFigures for "RNAIndel: discovering somatic coding indels from tumor RNA-Seq data"

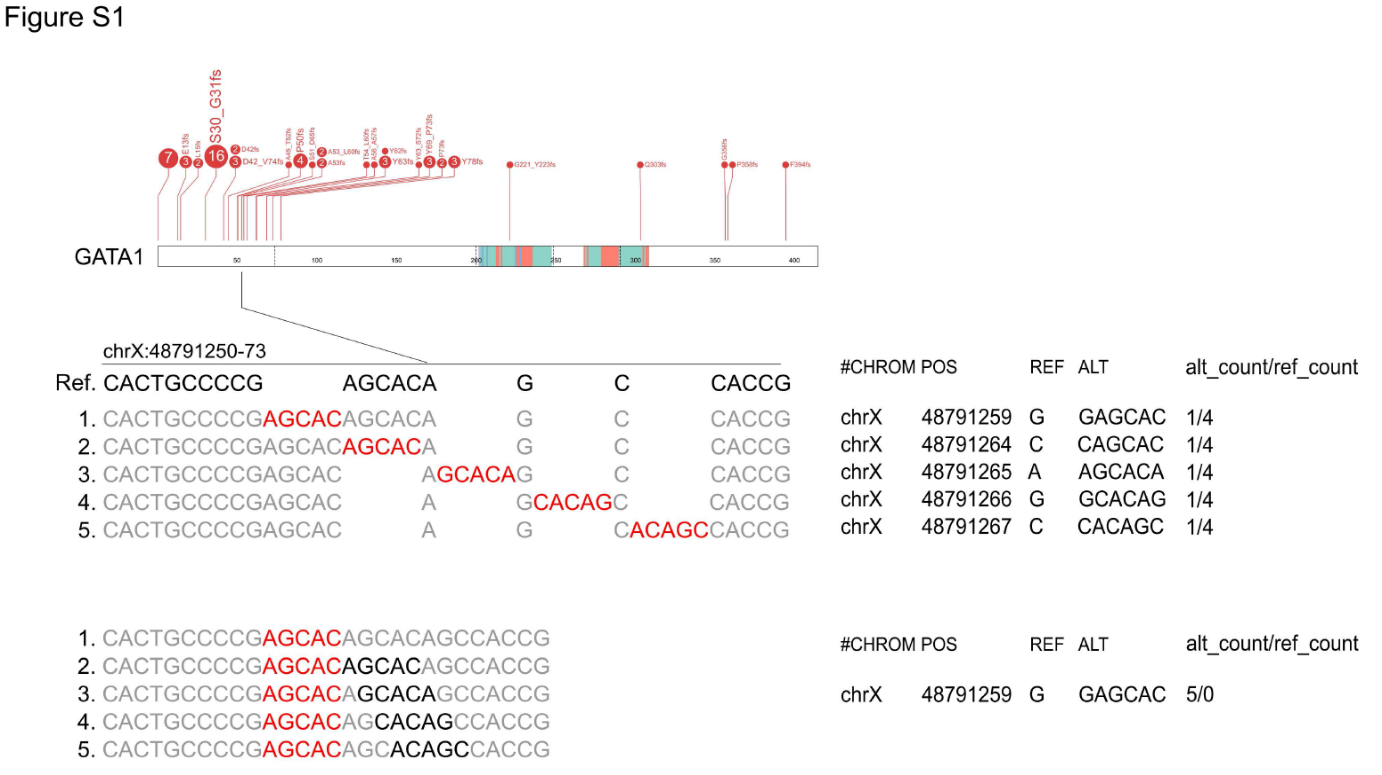

Figure S1. Equivalent indels. A 5-nt insertion in the *GATA1* gene that can be aligned to the reference in five different ways (*upper*). These alignments are left-aligned in red with the original inserted sequences highlighted in black (*lower*). Such alternative indel alignments are called equivalent and considered identical throughout the tool.

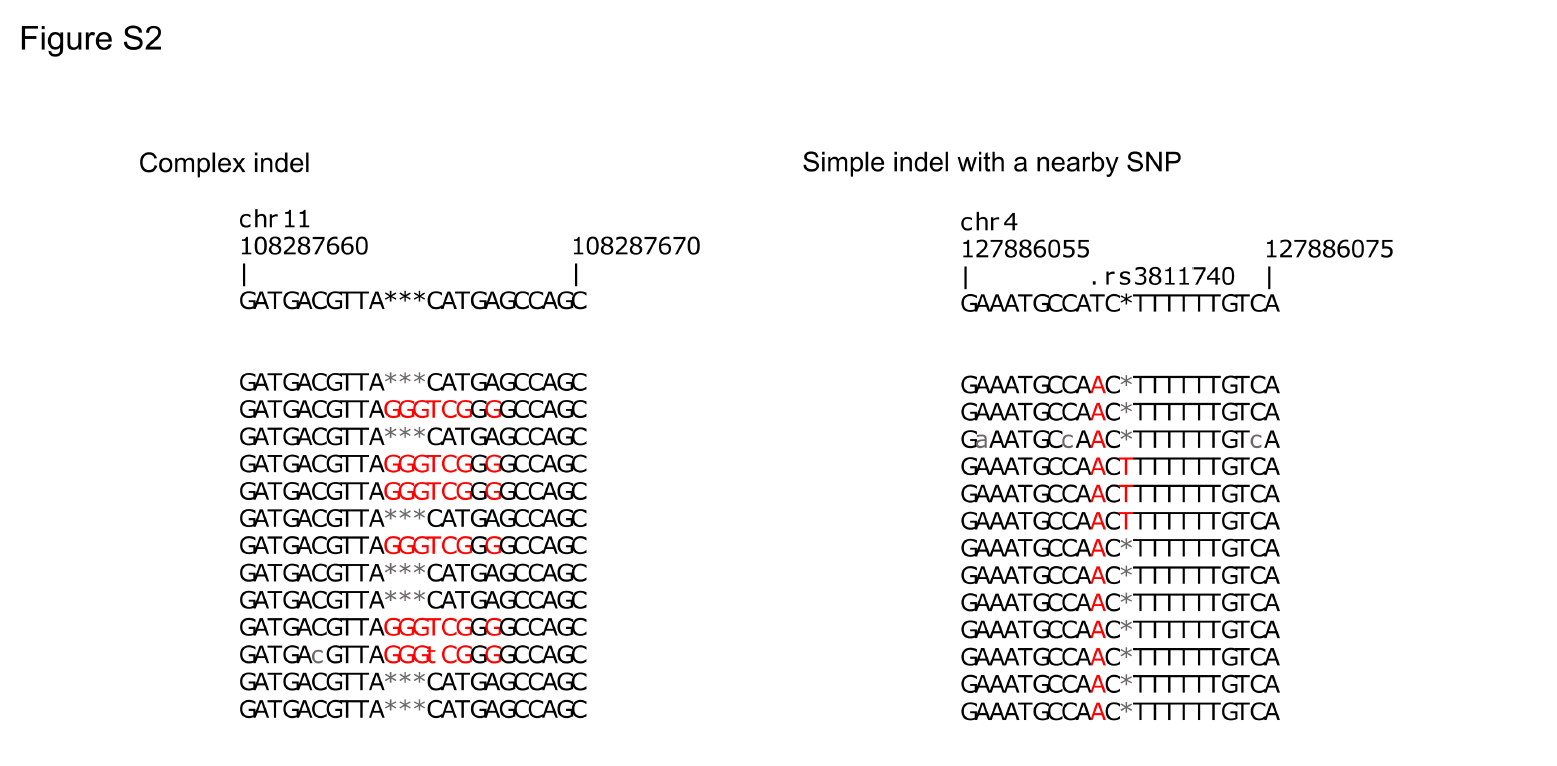

Figure S2. Complex indel. The inserted ‘GGGT’ is compounded with a cluster of misalignments (left). Note that the inserted sequence and additional misalignments are exclusively on the same reads. The inserted ‘T’ in the right is also associated with an additional misalignment (T>A), however this is a simple indel with a nearby single nucleotide polymorphism (rs3811740).

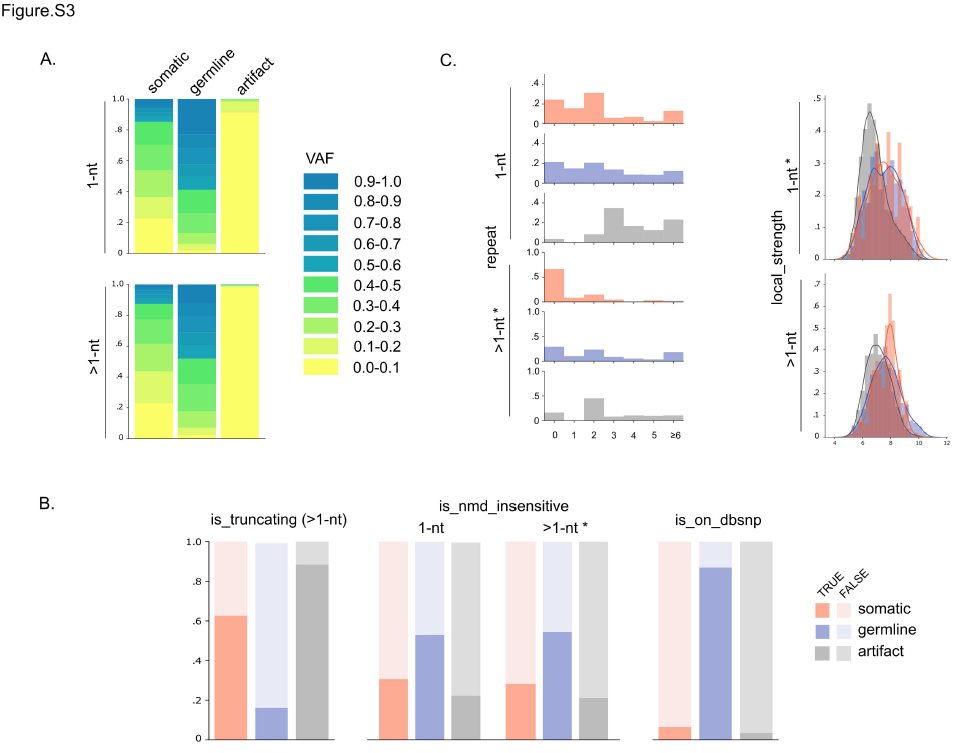

Figure S3. VAF spectrum, and germline and artifact features. **A**. The VAF spectrum of each class is visualized as a summary statistic of the features ‘ref_count’ and ‘alt_count’. **B**. Visualization of features characteristic of germline indels. The distribution of truncating indels was visualized for the feature ‘is_nmd_insensitive’. **C**. Visualization of features characteristic of artifacts. Asterisks indicate that the feature was shown for comparison, while not selected in the final model for the indel type (e.g., >1-nt * in ‘is_nmd_insensitive’).
